# ppx: Programmatic access to proteomics data repositories

**DOI:** 10.1101/2021.05.29.446304

**Authors:** William E Fondrie, Wout Bittremieux, William S Noble

## Abstract

The volume of proteomics and mass spectrometry data available in public repositories continues to grow at a rapid pace as more researchers embrace open science practices. Open access to the data behind scientific discoveries has become critical to validate published findings and develop new computational tools. Here, we present ppx, a Python package that provides easy, programmatic access to the data stored in ProteomeXchange repositories, such as PRIDE and MassIVE. The ppx package can either be used as a command line tool or a Python package to retrieve the files and metadata associated with a project when provided its identifier. To demonstrate how ppx enhances reproducible research, we used ppx within a Snakemake workflow to reanalyze a published dataset with the open modification search tool ANN-SoLo and compared our reanalysis to the original results. We show that ppx readily integrates into workflows and our reanalysis produced results consistent with the original analysis. We envision that ppx will be a valuable tool for creating reproducible analyses, providing tool developers easy access to data for development, testing, and benchmarking, and enabling the use of mass spectrometry data in data-intensive analyses. The ppx package is freely available and open source under the MIT license at: https://github.com/wfondrie/ppx

## Introduction

Open access to the data underlying published research is fundamental to foster transparency of the scientific process and to maximize the value of research for the public good. When the data is shared in a findable, accessible, interoperable, and reusable (FAIR) manner, it promotes reproducible research practices and enhances the reliability of the published findings [1, 2]. Data shared under FAIR principles may also yield biological insights beyond the original publication upon secondary analysis with alternative methods. Furthermore, open data provides an invaluable resource for tool developers to refine, test, and benchmark their tools, which ultimately benefits the field.

Fortunately, the proteomics and broader mass spectrometry communities have embraced data sharing and now consistently deposit the raw data behind their publications into public repositories [3]. The ProteomeXchange consortium [4] was formed in 2011 to coordinate the submission and dissemination of mass spectrometry proteomics data worldwide and has seen tremendous growth in the amount of data deposited in its partner repositories—PRIDE [5], PeptideAtlas [6], PASSEL [7], MassIVE [8], jPOST [9], iProx [10], and Panorama Public [11]—since its inception [12]. This push toward open data practices has proven immensely beneficial to the proteomics field, providing consistent datasets for tool development and benchmarking [13–17]. Additionally, the growing abundance of accessible data has been critical for the tools and insights that can only be gained from “big data,” including resources such as MassIVEKB [8] and PRIDE Cluster [18].

Easy, consistent access to the data in these repositories is critical to ensuring the reproducibility of proteomics analyses and leveraging the abundance of data to develop innovative computational approaches. However, much of this work is currently performed manually; often the task of downloading data is completed by navigating through the repository’s web interface to find the file transfer protocol (FTP) link for the specific project of interest. An exception to this pattern is the programmatic access to ProteomeXchange provided by the rpx R package [19].

Here, we present ppx, which provides a simple interface to proteomics repositories both as a command line tool and a Python package. Inspired by rpx, we developed ppx with the goal of improving the reproducibility of proteomics research and offering developers programmatic access to the growing tide of open mass spectrometry data.

## Methods and Results

### Installation and code availability

The ppx package is available for Python 3.6+ and can be easily installed from the Python Package Index (PyPI) with pip or via conda using the Bioconda channel [20]. The ppx package depends on the requests and tqdm [21] Python packages. As an open source project, ppx is publicly available on GitHub under the permissive MIT license: https://github.com/wfondrie/ppx.

### ppx design and implementation

The ppx package is a lightweight Python package that provides a consistent application programming interface (API) to ProteomeXchange and its partner repositories—currently PRIDE and MassIVE—with the goal to enable easy access to the files and metadata associated with each project. When users provide ppx with a ProteomeXchange, PRIDE, or MassIVE identifier, ppx is able to find the partner repository in which the project resides, retrieve metadata about the project, list the files associated with the project, and download the requested files to the user’s machine. To accomplish these tasks ppx leverages the metadata provided by the ProteomeXchange XML announcement for a project, as well as the metadata provided by PRIDE or MassIVE. In the case of PRIDE, we specifically use the RESTful API [22] to access information about a project.

We designed ppx to be most effective when project data is stored in a central directory on the user’s machine, which is the customizable default for ppx. When a resource is requested, ppx first checks whether the file or metadata has already been downloaded. By default, the remote repositories are only accessed when a new resource is requested. Furthermore ppx adopts a “lazy” approach to fetching data; remote resources are not accessed until they are needed. This behavior ensures that the repository servers are not unnecessarily burdened by requests, allows ppx to remain useful offline, and makes subsequent use of the same resources much faster for the user.

Downloading data via the ppx command line interface is easy. For example, a user can download the FASTA file from PXD000001 [23] using the following command:

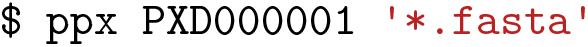

The ppx Python API offers greater flexibility and can be used to accomplish the same task:

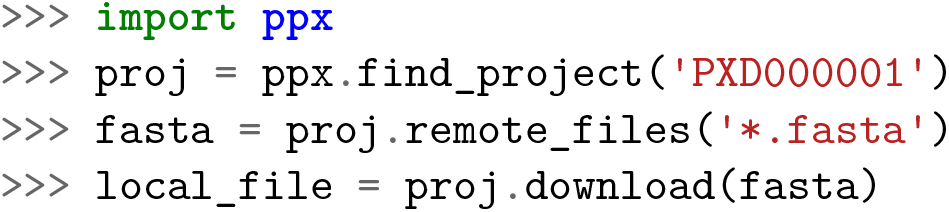

Thus, the files associated with a project are readily accessible from either command line interface or within a user’s Python session. Additionally, the ppx Python API provides methods to retrieve the paths to downloaded files and fetch project metadata, making it a powerful tool for data-intensive applications.

### ppx is a tool for reproducible research

When attempting to reproduce the results of a published proteomics study, often the first step is to download the raw mass spectrometry data and analyze it in the same manner as the original authors. This has historically been fraught with challenges—insufficient metadata, software installation issues, unpublished in-house code—which renders such exercises impossible. However, a recent trend in the proteomics community has been to embrace tools built for reproducible research. For example, the growth of resources such as Bioconda and BioContainers [25] have made it possible to manage software within virtual environments and containers, relieving the difficulties of installation and managing dependencies. The adoption of workflow engines such as Snakemake [26], Nextflow [27], and Cromwell [28] provide frameworks to run analysis pipelines reproducibly and orchestrate the necessary environments or containers. Furthermore, workflows built with these engines are readily transferred to the cloud, enabling them to scale massively [29]. Notably, ppx fits well within this framework by providing programmatic access to the data required to execute workflows.

As an example, we used ppx to help reanalyze a fractionated HEK293 cell line dataset [13] with the open modification spectral library search engine ANN-SoLo [24, 30]. Our goal was to reproduce the analysis originally presented by Bittremieux et al. [24], but with an updated version of ANN-SoLo. We used ppx to download the 24 mass spectrometry data files in Mascot generic format (MGF) and the spectral library directly from PRIDE project PXD009861. This library was the MassIVE-KB peptide spectral library (version 2017/11/27), concatenated with decoy spectra that were generated using the shuffle-and-reposition method [31], yielding 3,009,902 total spectra. We searched this data using ANN-SoLo version 0.3.3, as opposed to ANN-SoLo version 0.1.2 which was used for the original analysis. Notably, the newer version of ANN-SoLo supports the use of graphics processing units (GPUs) to massively speed up the search, at the cost of a slight loss statistical power to detect peptides [30]. Correspondingly, we chose search parameters that best matched the original ANN-SoLo analysis of the HEK293 data for our searches. Additionally, we set the hash length to 400 and used GPUs to accelerate our searches, leaving other parameters to their defaults.

After running our analysis, we compared the spectrum-spectrum matches (SSMs) detected at a 1% false discovery rate (FDR) in our reanalysis against the original results that were uploaded to the PRIDE project alongside the raw data. We found that the differences in precursor mass—or “mass shifts”—found in our reanalysis closely matched those that were originally reported (Figure 1A). As expected, we did observe a small loss of power in the total number of detected SSMs when using the GPU (Figure 1B); however, we also found that both the unmodified SSMs and those bearing the most common mass shifts were largely consistent between the analyses.

**Figure 1:**
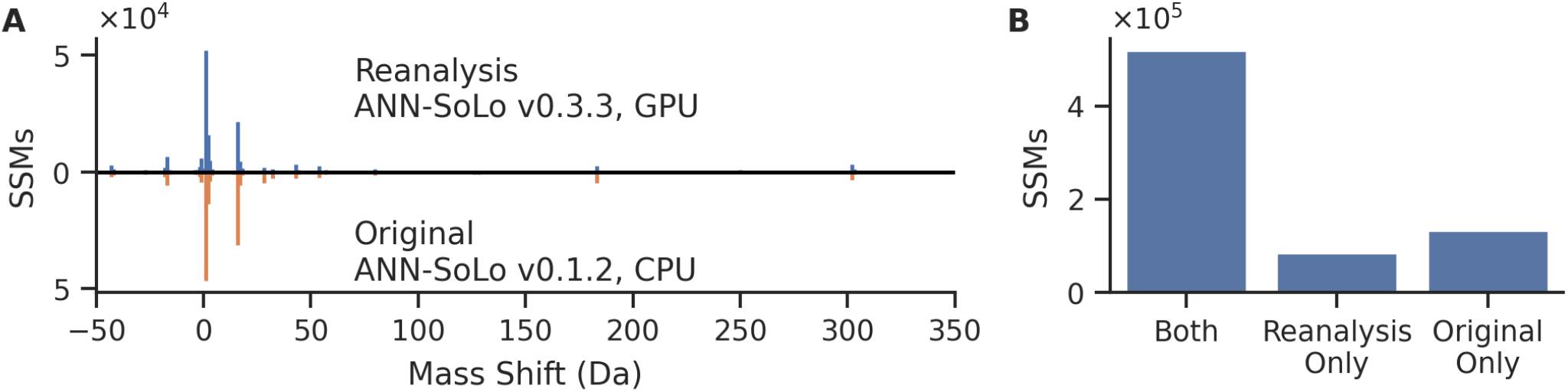
Reanalysis of the Chick et al. [13] HEK293 data with ANN-SoLo. (A) Our reanalysis using ANN-SoLo version 0.3.3 found similar mass shifts for SSMs accepted at 1% FDR when compared to the original analysis [24] conducted with ANN-SoLo version 0.1.2. (B) Although we observed some loss of power, a vast majority of the SSMs from the original analysis were recovered in our reanalysis.

Critically, every step of our reanalysis— downloading the raw data from PRIDE with ppx, searching with ANN-SoLo, downloading the original analysis results from PRIDE with ppx, and the creation of our figures with Python—is fully encapsulated in a Snakemake workflow that is publicly available at https://github.com/Noble-Lab/ppx-workflow. We chose to parallelize this analysis on our high-performance computing cluster, equipped with 12 NVIDIA RTX 2080 GPUs. However, the same workflow can be readily configured to reproduce our results on a single machine or in the cloud, provided sufficient computational resources are available.

## Conclusions

Here, we have introduced ppx and provided a short vignette on how ppx can be used in combination with a workflow engine to create fully reproducible analyses. The ppx package is a powerful tool for enabling reproducible research, and we envision that it will be particularly useful for tool developers to access the mountains of public mass spectrometry data at their disposal. Furthermore, the easy programmatic access to data that ppx provides will present great opportunities for the development of innovative approaches that require big data to be effective. Finally, we anticipate that ppx will become more valuable as more mass spectrometry experiments are annotated with standardized metadata [32], which lays the groundwork for fully automated analysis pipelines.

## Acknowledgments

The research reported in this publication was supported by the National Institutes of Health awards T32HG000035, P41GM103533, and R01GM121818. W.B. is a postdoctoral researcher of the Research Foundation – Flanders (FWO 12W0418N).

